# Dioxin-elicited decrease in cobalamin redirects hepatic propionyl-CoA metabolism to the β–oxidation-like pathway resulting in acrylyl-CoA conjugate accumulation

**DOI:** 10.1101/2021.03.24.436837

**Authors:** Karina Orlowska, Russ R. Fling, Rance Nault, Warren J. Sink, Anthony L. Schilmiller, Tim Zacharewski

## Abstract

2,3,7,8-Tetrachlorodibenzo-*p*-dioxin (TCDD) is a persistent environmental contaminant which induces diverse biological and toxic effects, including the reprograming of intermediate metabolism, mediated by the aryl hydrocarbon receptor (AHR). Targeted LC-MS analysis of hepatic extracts from mice gavaged with TCDD every 4 days for 28 days detected an increase in S-(2-carboxyethyl)-L-cysteine, a conjugate produced following the spontaneous reaction between the sulfhydryl group of cysteine and highly reactive acrylyl-CoA, an intermediate in the cobalamin (Cbl)-independent β–oxidation-like metabolism of propionyl-CoA. In addition to repressing genes in both the canonical Cbl-dependent carboxylase and the alternate Cbl-independent β–oxidation-like pathways at 30 μg/kg TCDD, methylmalonyl-CoA mutase (MUT) activity was inhibited at lower doses. Moreover, TCDD decreased serum Cbl levels and hepatic cobalt levels while eliciting negligible effects on gene expression associated with Cbl absorption, transport, trafficking, or the derivatization to 5’-deoxy-adenosylcobalamin (AdoCbl), the required MUT cofactor. In addition to inducing *Acod1* that encodes for aconitate decarboxylase 1, the enzyme responsible for the decarboxylation cis-aconitate to itaconate, TCDD also dose-dependently increased itaconate levels in hepatic extracts. MUT inhibition is consistent with itaconate activation to itaconyl-CoA, a MUT suicide inactivator that adducts AdoCbl, that in turn, inhibits MUT activity and reduces Cbl levels. Collectively, these results suggest the decrease in MUT activity is due to Cbl depletion following TCDD treatment that redirected propionyl-CoA metabolism to the alternate Cbl-independent β–oxidation-like pathway. The resulting hepatic accumulation of acrylyl-CoA likely contributes to TCDD-elicited hepatotoxicity and the multi-hit progression of steatosis to steatohepatitis with fibrosis.

## Introduction

Adverse effects elicited by exposure to toxic substances are not only influenced by the dose, route of administration, and exposure duration but also genetic, epigenetic and other systemic factors. Cellular responses to xenobiotic insults are essential to minimize damage and ensure survival. Adaptive effects, such as cytochrome P450 induction metabolize and facilitate xenobiotic detoxification and excretion, while immune cell infiltration expedites damaged cell removal with metabolic reprogramming supporting increased glutathione biosynthesis. This culminates in an array of apical responses that may lead to inflammation, repair, proliferation and/or additional cytotoxicity. Xenobiotics may also trigger the differential expression of genes and modulate enzyme activities that have the potential to qualitatively and quantitatively alter endogenous metabolite profiles with the possibility of mitigating or exacerbating the overall toxic burden. Elucidating the role of endogenous metabolic plasticity in response to foreign agents, whether drugs, environmental contaminants or natural products, is essential to the identification of susceptible cell sub-types, and to distinguish adverse from adaptive responses underlying toxic effects. In addition to discovering potential strategies to reduce off-target toxicity, elucidating the mechanisms involved may reveal novel vulnerabilities for exploitation as innovative therapeutic approaches for the treatment of diseases with similar pathologies.

The progression of simple, reversible hepatic fat accumulation to steatohepatitis with fibrosis and hepatocyte ballooning describes the typical clinicopathologic spectrum of phenotypes associated with non-alcoholic fatty liver disease (NAFLD). In NAFLD, >5% of the cytosolic space within hepatocytes is occupied by lipid droplets in patients where little to no alcohol was consumed and there was no secondary cause involving viral hepatitis, medication, or lipodystrophy (1). The ‘two-hit’ hypothesis for NAFLD development has evolved into a multiple hit etiology that disrupts several pathways (2). NAFLD prevalence is projected to increase from ~83 million in 2015 to ~101 million by 2030 in the US alone, while increasing the risk for more complex disorders including Metabolic Syndrome, cardiovascular disease, diabetes, cirrhosis, end-stage liver disease and hepatocellular carcinoma (HCC) (3, 4). Furthermore, progression of NAFLD to non-alcoholic steatohepatitis (NASH) is the leading indication for liver transplantation and the third leading cause of hepatocellular carcinoma in the US with limited treatment options (5–7).

Diet, lifestyle, and genetic background are known factors that contribute to NAFLD development and progression. Environmental contaminants also induce steatosis, suggesting a possible role in disease etiology (8, 9). For example, several pesticides, solvents and their metabolites induce hepatic fat accumulation with 2,3,7,8-tetrachlorodibenzo-*p*-dioxin (TCDD) and related compounds exhibiting the greatest potency (10). In mice, TCDD dose-dependently induced micro- and macro-steatosis with marked increases in hepatic unsaturated fatty acids (FAs), triacylglycerols (TAGs), phospholipids and cholesterol esters (CEs) (11–14). This has been attributed to increased hepatic uptake of dietary and mobilized peripheral fats, reduced very low density lipoprotein (VLDL) export, and the inhibition of hepatic FA oxidation (15, 16). In humans, TCDD and related compounds have been associated with dyslipidemia and inflammation (17–20). Epidemiological studies also report elevated serum cholesterol and TAG levels in exposed workers (21–24), while *in utero* TCDD exposure increased the risk for Metabolic Syndrome in male offspring (25).

TCDD is the prototypical member of a class of persistent environmental contaminants that includes polychlorinated dibenzodioxins (PCDDs), dibenzofurans (PCDFs) and biphenyls (PCBs). Congeners with lateral chlorines induce a plethora of species-, sex-, tissue-, and cell-specific responses (26). TCDD and co-planar PCBs are classified as IARC group 1 human carcinogens while evidence for the carcinogenicity of other toxic PCDDs and PCDFs remains equivocal (27, 28). TCDD and related compounds are non-genotoxic and most, if not all, of their effects are mediated by the aryl hydrocarbon receptor (AhR), a ligand activated basic helix-loop-helix PER-ARNT-SIM transcription factor. Although a number of structurally diverse chemicals, endogenous metabolites, microbial products and natural products activate the AhR, its physiological ligand is unknown. Following ligand binding and the dissociation of chaperone proteins, the activated AhR translocates from the cytosol to the nucleus and dimerizes with the AhR nuclear translocator (ARNT). This heterodimer then binds dioxin response elements (DREs; 5’-GCGTG-3’) as well as non-consensus sites throughout the genome recruiting coactivator complexes to gene promoters to elicit differential expression (29). AHR-mediated toxicity is generally believed to be the result of dysregulated gene expression. However, the consequences of AhR-mediated differential gene expression and the associated effects on intermediate metabolism of endogenous metabolites have not been explored.

The emergence of transcriptomics and metabolomics has provided tools to comprehensively assess the time- and dose-dependent impacts of drugs, chemicals, environmental contaminants, and natural products on gene expression and intermediate metabolism as well as resulting pathologies. Several studies have reported the effects of PCDDs, PCDFs or PCBs on gene expression and/or endogenous metabolite levels in diverse *in vivo* and *in vitro* models (11, 12, 36, 13, 14, 30–35). In this study, we tested the hypothesis that the dose-dependent disruption of propionyl-CoA metabolism produces toxic intermediates that contribute to TCDD hepatotoxicity and progression of steatosis to steatohepatitis with fibrosis. Our results suggest TCDD dose-dependently reduced cobalamin (Cbl aka vitamin B12) levels compromising methylmalonyl-CoA mutase (MUT) activity and limiting the metabolism of propionyl-CoA to succinyl-CoA using the canonical Cbl-dependent carboxylation pathway. Consequently, accumulating propionyl-CoA was redirected to the alternate Cbl-independent β–oxidation-like pathway resulting in the dose-dependent accumulation of acrylyl-CoA, as indicated by the increase in S-(2-carboxyethyl)-L-cysteine (SCEC), a conjugate produced following the spontaneous reaction between the sulfhydryl group of cysteine and highly reactive acrylyl-CoA.

## MATERIALS AND METHODS

### Animal Treatment

Postnatal day 25 (PND25) male C57BL/6 mice weighing within 10% of each other were obtained from Charles River Laboratories (Kingston, NY). Mice were housed in Innovive Innocages (San Diego, CA) containing ALPHA-dri bedding (Shepherd Specialty Papers, Chicago, IL) in a 23°C environment with 30– 40% humidity and a 12 hr/12 hr light/dark cycle. Aquavive water (Innovive) and Harlan Teklad 22/5 Rodent Diet 8940 (Madison, WI) containing 60 μg vitamin B_12_/kg of diet was provided *ad libitum*. On PND28, mice were orally gavaged at the start of the light cycle (zeitgeber [ZT] 0-1) with 0.1 ml sesame oil vehicle (Sigma-Aldrich, St. Louis, MO) or 0.01, 0.03, 0.1, 0.3, 1, 3, 10, and 30 μg/kg body weight TCDD (AccuStandard, New Haven, CT) every 4 days for 28 days for a total of 7 treatments. The first gavage was administered on day 0, with the last gavage administered on day 24 of the 28-day study. The doses used consider the relatively short study duration in mice compared to lifelong cumulative human exposure from diverse AhR ligands, the bioaccumulative nature of halogenated AhR ligands, and differences in the half-life of TCDD between humans (1–11 years (37, 38)) and mice (8–12d (39)). Similar treatment regimens have been used in previous studies (11, 16, 31, 35, 40). On day 28, tissue samples were harvested (ZT 0-3), immediately flash frozen in liquid nitrogen and stored at −80°C until analysis. All animal procedures were in accordance with the Michigan State University (MSU) Institutional Animal Care and Use Committee ethical guidelines and regulations.

### Liquid Chromatography Tandem Mass Spectrometry (LC-MS)

Flash frozen liver samples (~25 mg) were extracted using HPLC-grade methanol and water (5:3 ratio) containing 20 ^13^C-,^15^N-labelled amino acid (Sigma; 767964) internal standards (31). HPLC-grade chloroform (methanol:water:chloroform ratio 5:3:5) was added, vortexed, shaken for 15 min at 4 °C, and centrifuged at maximum speed (3000 × g) to achieve phase separation. The methanol:water phase containing the polar metabolites was transferred, dried under nitrogen gas at room temperature. Untargeted extractions were reconstituted with 400 μl of 10 mM tributylamine and 15 mM acetic acid in 97:3 water:methanol for analysis. Samples were analyzed on a Xevo G2-XS Quadrupole Time of Flight (QTof) mass spectrometer attached to a Waters Acquity UPLC (Waters, Milford, MA) with negative-mode electrospray ionization run using an MS^E^ continuum mode method. LC phases, gradient rates, and columns were used as previously published (31). For untargeted acyl-CoA analysis, MS^E^ continuum data was processed with Progenesis QI (Waters) to align features, deconvolute peaks, and identify metabolites. Metabolite identifications were scored based on a mass error <12 ppm to Human Metabolome Database entries (41), isotopic distribution similarity, and theoretical fragmentation comparisons to MS^E^ high-energy mass spectra using the MetFrag option. Raw signals for each compound abundance were normalized to a correction factor calculated using the Progenesis QI median and mean absolute deviation approach. Significance was determined by a one-way ANOVA adjusted for multiple comparisons with a Dunnett’s *post-hoc* test. Raw data is deposited in the NIH Metabolomics Workbench (ST001379).

For S-(2-carboxyethyl)-L-cysteine (SCES) analysis, frozen liver samples (~25 mg) were homogenized in 400 μl of 70% acetonitrile containing 2 μM ^13^C_5_,^15^N-methionine (labeled amino acid cocktail; Sigma). After centrifugation, the supernatant was transferred on Resprep PLR 96-well plate (Restek) where precipitated proteins and phospholipids were removed by filtration. Filtrate was dried under nitrogen gas and reconstituted in 10 mM perfluoroheptanoic acid (PFHA) in water. Samples were analyzed using a Xevo TQ-S micro Triple Quadrupole Mass Spectrometry system combined with Waters Acquity UPLC (Waters, Milford, MA) fitted with an Acquity HSS T3 2.1 x 100 mm column maintained at 40°C. The mobile phases were 10mM PFHA in water (mobile phase A) and acetonitrile (mobile phase B) using the following gradient: 0 min – 100% A, 1.0 min – 100% A, 6.0 min – 35% A, 6.01 min – 10% A, 7.0 min – 10% A, 7.01 min – 100% A, 9.0 min – 100% A). The flow rate was 0.3 mL/min with an injection volume of 10 μL and total run time 9 minutes. Multiple reaction monitoring in positive ion mode was used for compound detection. Transitions used for quantification are provided in Supplementary Table S1.

Itaconic acid extracts were prepared and analyzed as previously described with slight modifications (42). Briefly, frozen liver samples (~40mg) were homogenized (Polytron PT2100, Kinematica, Lucerne, Switzerland) in acetonitrile:water ratio 8:2, vortexed, shaken for 5 min at 4 °C, and centrifuged at maximum speed (3000×g). Supernatant was dried under nitrogen and dried extracts were reconstituted in 200 μL of a 1:9 methanol:water solution containing 2% formic acid. Samples were separated with an Acquity HSS T3 column (1.8 μm, 100 × 2.1 mm; Waters, Milford, MA) with 0.1% formic acid in water (solvent A) and 0.1% formic acid in acetonitrile (solvent B) mobile phase gradient (0 min – 100% A, 1.0 min – 100% A, 2.0 min – 80% A, 4.0 min – 1% A, 5.0 min – 1% A, 5.01 min – 100% A, 7.0 min – 100% A, flow rate 0.3 mL/min). Detection was performed using a Xevo G2-XS QTof mass spectrometer with electrospray ionization in negative ion mode. Data were acquired using a TOF MS scanning method (m/z 50-1200 scan range) with the target enhancement option tuned for m/z 129. Signals were identified by retention time and accurate mass using MassLynx Version 4.2 (Waters). A six point itaconic acid standard calibration curve was prepared by diluting unlabeled standard (Sigma Aldrich).

### Gene Expression, ChIP, pDRE and Protein Location Data

Hepatic RNA-seq data sets were previously published (31, 32, 43). Genes were considered differentially expressed when |fold-change| ≥ 1.5 and posterior probability values (P1(*t*)) ≥ 0.8 as determined by an empirical Bayes approach (44). Hepatic time course (GSE109863), dose response (GSE203302), and diurnal rhythmicity (GSE119780) datasets as well as duodenal (GSE87542), jejunal (GSE90097), proximal (GSE171942) and distal ileal (GSE89430) and colon (GSE171941) datasets are available at the Gene Expression Omnibus. Diurnal rhythmicity was determined using JTK_CYCLE as previously described (31). AhR ChIP-seq (GSE97634) and computationally identified putative dioxin response elements (pDREs, https://doi.org/10.7910/DVN/JASCVZ) data were previously published (32, 45). Significant AhR ChIP-seq binding used a false discovery rate (FDR) ≤ 0.05. pDREs were considered functional with a matrix similarity score (MSS) ≥ 0.856.

### Quantitative Real-Time Polymerase Chain Reaction (qRT-PCR)

Expression of *Acod1* was determined by qRT-PCR. Total hepatic RNA was reverse transcribed by SuperScript II (Invitrogen, Carlsbad, CA) using oligo dT primer according to the manufacturer’s protocol. The qRT-PCR was performed using iQ SYBR Green Supermix (BioRad, Hercules, CA) on a Bio-Rad CFX Connect Real-Time PCR Detection System. Gene expression relative to vehicle control was calculated using the 2^-ΔΔCT^ method, where each sample was normalized to the geometric mean of 3 housekeeping genes (*Actb, Gapdh*, and *Hprt*). Gene expression data are plotted relative to vehicle control. See Supplementary Table S3 for primer sequences.

### Measurement of Cbl and Cobalt Levels

Serum Cbl levels (vehicle, 1–30 μg/kg TCDD groups) were determined by ELISA using a commercially available kit (Cusabio, Houston, TX) using SpectraMax ABS Plus plate reader (Molecular Devices, San Jose, CA). Cobalt levels were measured in liver extracts (vehicle, 3–30 μg/kg TCDD groups) using inductively coupled plasma mass spectrometry (ICP-MS) at the MSU Diagnostic Center for Population and Animal Health (DCPAH).

### MUT assay

MUT activity was measure in hepatic extracts using a thiokinase-coupled, spectrophotometric assay (46, 47). Total protein lysates were isolated from frozen samples with NP-40 cell lysis buffer (Thermo Fisher Scientific, Waltham, MA) containing protease inhibitor using a Polytron PT2100 homogenizer (Kinematica). MUT activity assay mixture contained 50 mM trisphosphate buffer pH 7.5, 4 mM 5,5’-dithiobis-(2-nitrobenzoic acid) (DTNB), 10 mM ADP, 20 mM MgCl_2_, 20 mM methylmalonyl-CoA, 0.1 U thiokinase, 20 μM AdoCbl. All components were incubated at 30°C for 40 min to equilibrate temperature. The reaction was started by the addition of 1 μg protein extract. Absorbance at 412 nm (A412) was measured for 2 min, every 8 seconds. The increase in A412 in reactions lacking substrate was subtracted from all readings. A412 values were converted to concentration of free CoA using a pathlength correction determined for the reaction volume and extinction coefficient of 14150 M^-1^ cm^-1^.

### Metagenomic Analysis of Microbial Cbl Metabolism

Cecums from vehicle, 0.3, 3, and 30 μg/kg TCDD treatment groups were used for metagenomic analysis. Genomic DNA was extracted using the FastDNA spin kit for soil (SKU 116560200, MP Biomedicals, Santa Ana, CA) and submitted for quality control, library prep and 150-bp paired-end sequencing at a depth ≥136 million reads using an Illumina NovaSeq 6000 (Novogene, Sacramento, CA) (NCBI BioProject ID: PRJNA719224). Reads aligning to the C57BL/6 *Mus musculus* genome (NCBI genome assembly: GRCm38.p6) were identified, flagged and removed using bowtie2 (48), SamTools (49) and bedtools (50). The HuMaNn3 bioinformatic pipeline (51) was used with default settings to classify reads to UniRef90 protein ID’s using UniProt’s UniRef90 protein data base (January, 2019). Reads classified to UniRef90 IDs were mapped to enzyme commission (EC) and PFAM entries using the human_regroup_table tool. Abundances were normalized to gene copies per million reads using the human_renorm_table tool. Statistical analysis used Maaslin2 (https://github.com/biobakery/Maaslin2) with default settings for normalization (total sum scaling), analysis method (general linear model), and multiple correction adjustment.

## RESULTS

### LC-MS/MS analysis

Untargeted metabolomics annotation suggested TCDD-elicited dose-dependent changes in the level of intermediates associated with the Cbl-independent β-oxidation-like metabolism of propionyl-CoA. More specifically, this included the presence of acrylyl-CoA and 3-hydroxypropionyl-CoA in hepatic extracts following oral gavage with TCDD every 4 days for 28 days (data available at NIH Metabolomics Workbench, ST001379). However, the untargeted annotations were not sufficient to conclude that propionyl-CoA was metabolized via the Cbl-independent β–oxidation-like pathway (Fig 2A). Targeted analysis was unsuccessful in confirming the identity of acrylyl-CoA due its high reactivity. Alternatively, the presence of acrylyl-CoA was confirmed following the detection of S-(2-carboxyethyl) cysteine (SCEC), a conjugate formed following the spontaneous reaction between acrylyl-CoA and the sulfhydryl group of cysteine (52). Hepatic SCEC levels increased 15.6-fold following oral gavage of male mice with 30 μg/kg TCDD every 4 days for 28 days (**Table 1**). Urine cysteine/cysteamine conjugates with acrylyl-CoA are routinely used to diagnose Leigh syndrome where the conversion of acrylyl-CoA to 3-hydroxypropionyl-CoA is inhibited due to a deficiency in the short chain enoyl-CoA hydratase (ECHS1) activity (52). The increased levels of SCEC not only confirmed acrylyl-CoA accumulation but also ECHS1 inhibition, and that propionyl-CoA was metabolized via the alternate Cbl-independent β–oxidation-like pathway as opposed to the preferred Cbl-dependent carboxylation pathway (53).

**Figure 1.**
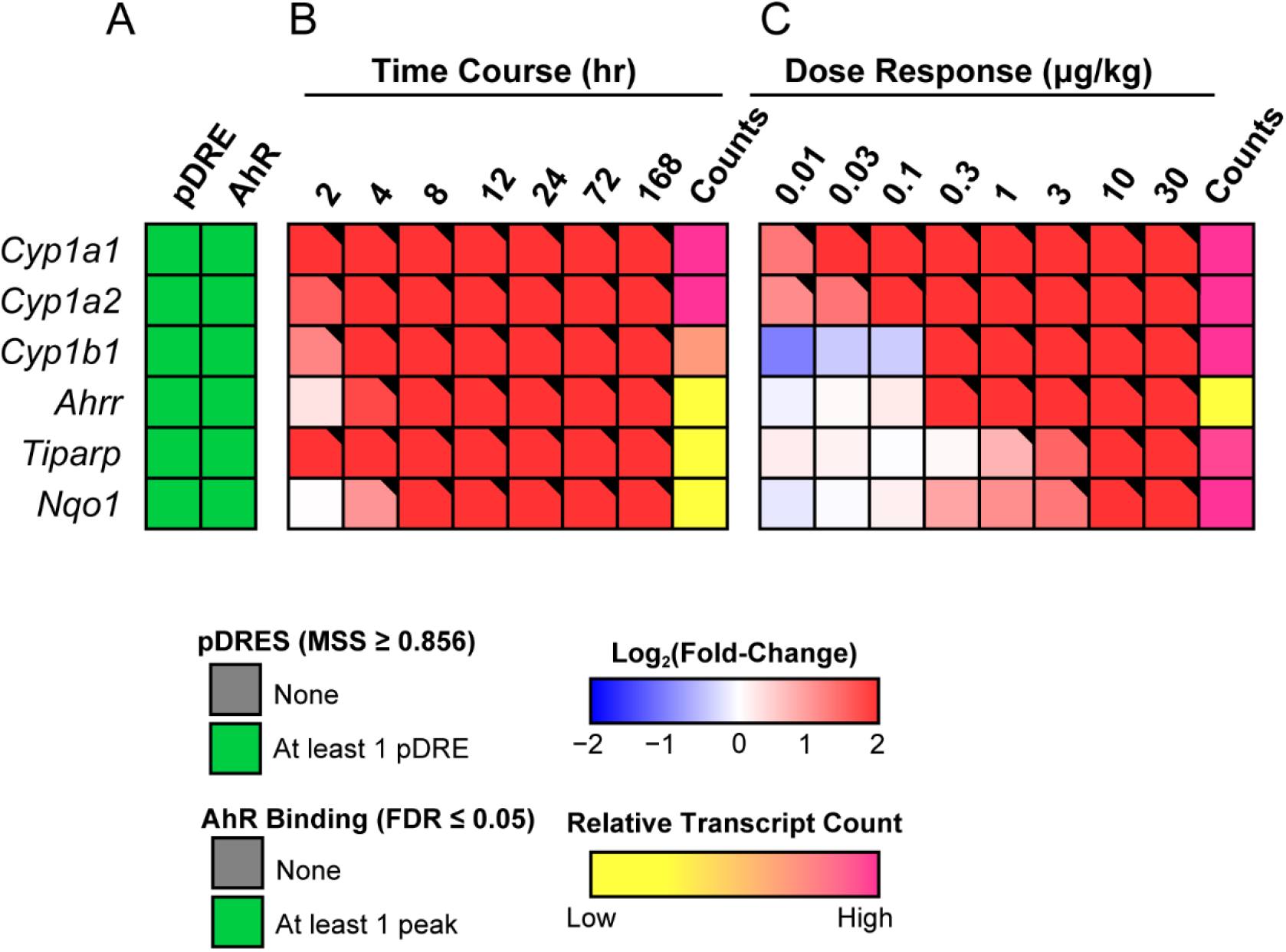
Effects of TCDD on the expression of AhR target genes. **A)** The presence of putative dioxin response elements (pDREs) and AhR genomic binding 2 hrs after a single bolus dose of 30 μg/kg TCDD. **B)** Hepatic expression of AhR target genes assessed in a time course study. Male C57BL/6 mice (n=3) were administered a single bolus dose of 30 μg/kg TCDD. Liver samples were collected at the corresponding time point. Color scale represents the log2(fold change) for differential gene expression determined by RNA-Seq analysis. Counts represent the maximum number of raw aligned reads for any treatment group. Low counts (<500 reads) are denoted in yellow with high counts (>10,000) in pink. **C)** Dose-dependent gene expression was assessed in mice (n=3) following oral gavage with sesame oil vehicle or TCDD. Differential expression with a posterior probability (P1(*t*))>0.80 is indicated with a black triangle in the upper right tile corner.

**Figure 2.**
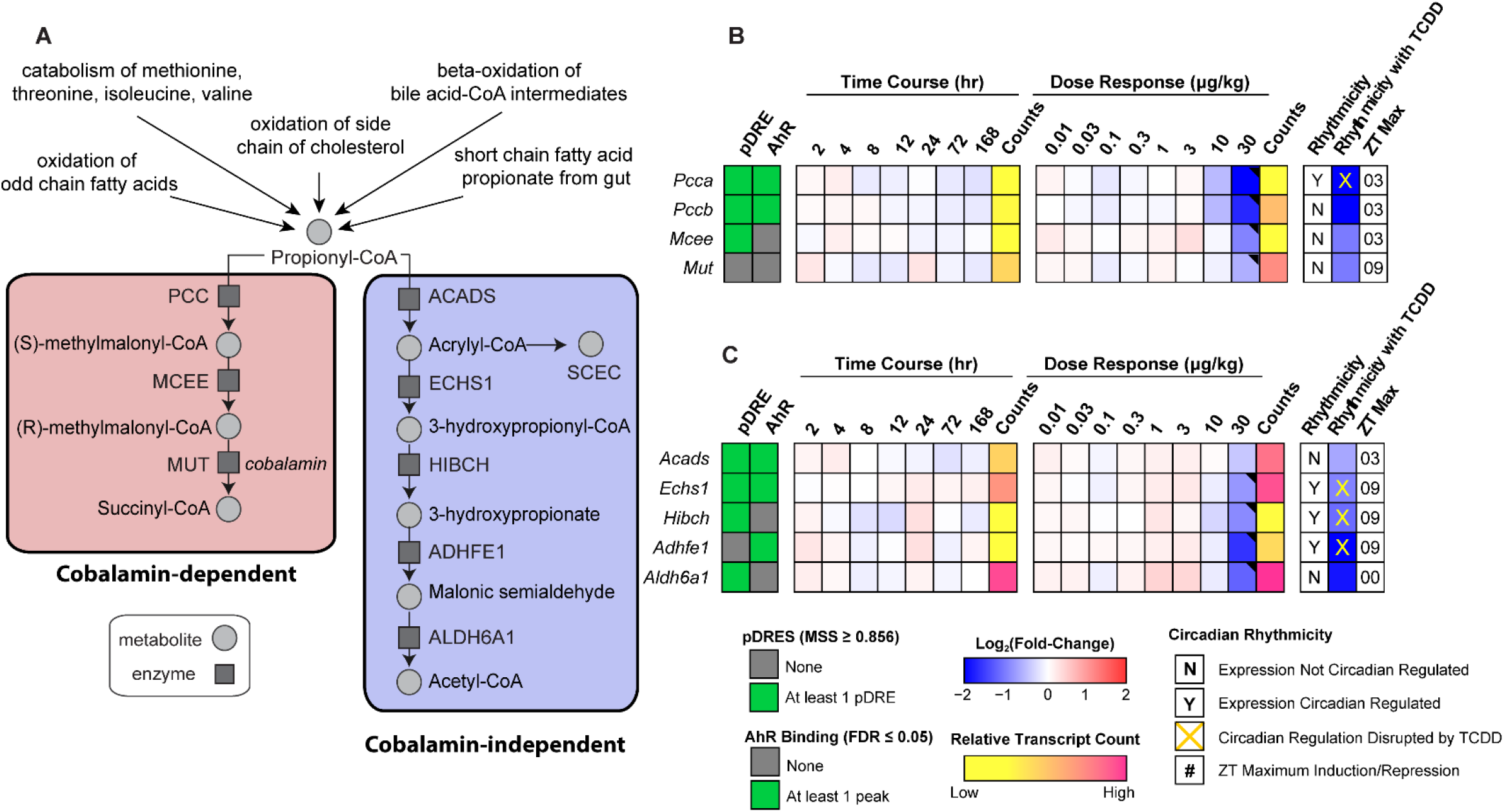
Effects of TCDD on the expression of genes associated with propionyl-CoA metabolism. **A)** Schematic pathway depicting enzymes and metabolites associated with propionyl-CoA metabolism via the cobalamin (Cbl)-dependent carboxylation pathway or the Cbl-independent β–oxidation-like pathway. **B)** Heatmap for genes associated with the propionyl-CoA canonical carboxylation pathway. **C)** Heatmap for genes associated with the Cbl-independent propionyl-CoA β-oxidation-like pathway. Hepatic expression of genes associated with propionyl-CoA was assessed in a time course- and dose-dependent manner. In time course study male C57BL/6 mice (n=3) were administered a single bolus dose of 30 μg/kg TCDD, after which tissue was collected at the corresponding timepoint, while in dose-dependent study male C57BL/6 mice (n=3) were orally gavaged with sesame oil vehicle or TCDD every 4 days for 28 days. The presence of putative dioxin response elements (pDREs) and AhR binding to the intragenic region represents as green boxes. Color scale represents the log2(fold change) for differential gene expression determined by RNA-Seq analysis. Counts represents the maximum raw number of aligned reads to each transcript where a lower level of expression (≤500 reads) is depicted in yellow and a higher level of expression (≥10,000) is depicted in pink. Genes that are circadian regulated are denoted by “Y”. Disruption of circadian rhythmicity following oral gavage with 30 μg/kg TCDD every 4 days for 28 days is denoted by an orange ‘X’. The ZT with maximum induction/repression is shown for each gene. Differential expression with a posterior probability (P1(*t*))>0.80 is indicated with a black triangle in the upper right tile corner.

**Table 1.** S-(2-carboxyethyl) cysteine fold changes in comparison to vehicle in liver extracts (n=5, ± S.E.M.) assessed by targeted liquid chromatography tandem mass spectrometry. Mice were orally gavaged every 4 days for 28 days with TCDD (or sesame oil vehicle). Asterisk (*) denotes significance (p≤0.05) determined by one-way ANOVA with Dunnett’s *post-hoc* testing.

| Compound name | Molecular mass | Retention Time (min) | Fold-Change (TCDD vs Vehicle) |  |  |  |  |
| --- | --- | --- | --- | --- | --- | --- | --- |
|  |  |  | 0.3 µg/kg | 1 µg/kg | 3 µg/kg | 10 µg/kg | 30 µg/kg |
| S-(2-carboxyethyl) cysteine | 194 | 2.6 | 1.10 ± 0.34 | 1.16 ± 0.12 | 1.16 ± 0.17 | 1.70 ± 0.42 | **15.62 ± 2.59\*** |

Note that the dose range and treatment regimen used in this study resulted in hepatic TCDD levels that approached steady state while inducing full dose response curves for known AhR target genes (**Fig. 1**) in the absence of (i) necrosis or apoptosis, (ii) marked increases in serum alanine transaminase (ALT), (iii) changes in food consumption and (iv) body weight loss >15% (31, 35). At 0.01 μg/kg, hepatic TCDD levels were comparable to control levels, and to background dioxin-like compound levels reported in US, German, Spanish and British serum samples (54) while mid-range doses yielded levels comparable to women from the Seveso Health Study (18). At 30 μg/kg TCDD, mouse hepatic tissue levels were comparable to serum levels reported in Viktor Yushchenko following intentional poisoning (37). Consequently, the metabolomics and gene expression effects elicited by TCDD, described below, cannot be attributed to overt toxicity.

### Propionyl-CoA metabolism gene expression effects

**Fig. 2** summarizes the temporal- and dose-dependent effects of TCDD on gene expression associated with propionyl-CoA metabolism. Note that all reported hepatic fold-changes discussed in the text of this study were taken from the circadian gene expression data set (GSE119780) at the optimal zeitgeber time (ZT0-3) unless otherwise indicated.

ChIP-seq analysis 2 hrs after a bolus oral gavage of 30 μg/kg TCDD suggested AhR enrichment may be involved in the repression of propionyl-CoA carboxylase (*Pcca* and *Pccb* subunits, 2.7- and 2.9-fold, respectively), short chain enoyl-CoA hydratase (*Echs1*, 1.5-fold) and alcohol dehydrogenase iron containing 1 (*Adhfe1*, 2.2-fold) but not for aldehyde dehydrogenase 6 family member A1 (*Aldh6a1*, 2.2-fold) (**Fig. 2B, C**). Moreover, the effects of TCDD on gene expression associated with both pathways were negligible within the 168 hrs time course study. TCDD disrupted the rhythmic expression of *Pcca, Echs1*, 3-hydroxyisobutyryl-CoA hydrolase (*Hibch*), and *Adhfe1*, all of which exhibited oscillating expression due to diurnal regulation. At 30 μg/kg, TCDD repressed *Pcca* and *Pccb* of the Cbl-dependent carboxylation pathway suggesting propionyl-CoA metabolism was redirected to the alternative Cbl-independent β–oxidation-like pathway. Although 30 μg/kg TCDD also repressed genes associated with the alternative pathway, the expression of short chain acyl-CoA dehydrogenase (*Acads*) was not affected by treatment, allowing propionyl-CoA to be oxidized to acrylyl-CoA. This suggested TCDD-elicited gene repression may contribute to the redirection of propionyl-CoA metabolism from the preferred Cbl-dependent carboxylation pathway to the alternate Cbl-independent β–oxidation-like pathway at the highest dose.

### Effects on Cbl and cobalt levels

Elevated levels of acrylyl-CoA and 3-hydroxypropionate (3-HP), as well as their derivatives, are not normally detected at appreciable levels in healthy individuals (55). Acrylyl-CoA and 3-HP typically accumulate following disruption of the canonical Cbl-dependent propionate catabolism pathway due to Cbl deficiency or mutations within propionyl-CoA carboxylase or MUT that affect enzyme activity (56). Since MUT is only one of two mammalian enzymes known to be Cbl dependent for activity, we examined the effects of TCDD on the levels of Cbl and cobalt, the metal ion that occupies the coordinate center of the corrin ring. TCDD dose-dependently reduced total serum Cbl levels and cobalt levels in hepatic extracts (**Fig. 3**). Therefore, reduced Cbl deficiency may be responsible for lower MUT activity and affect propionyl-CoA metabolism via the canonical Cbl-dependent carboxylation pathway.

**Figure 3.**
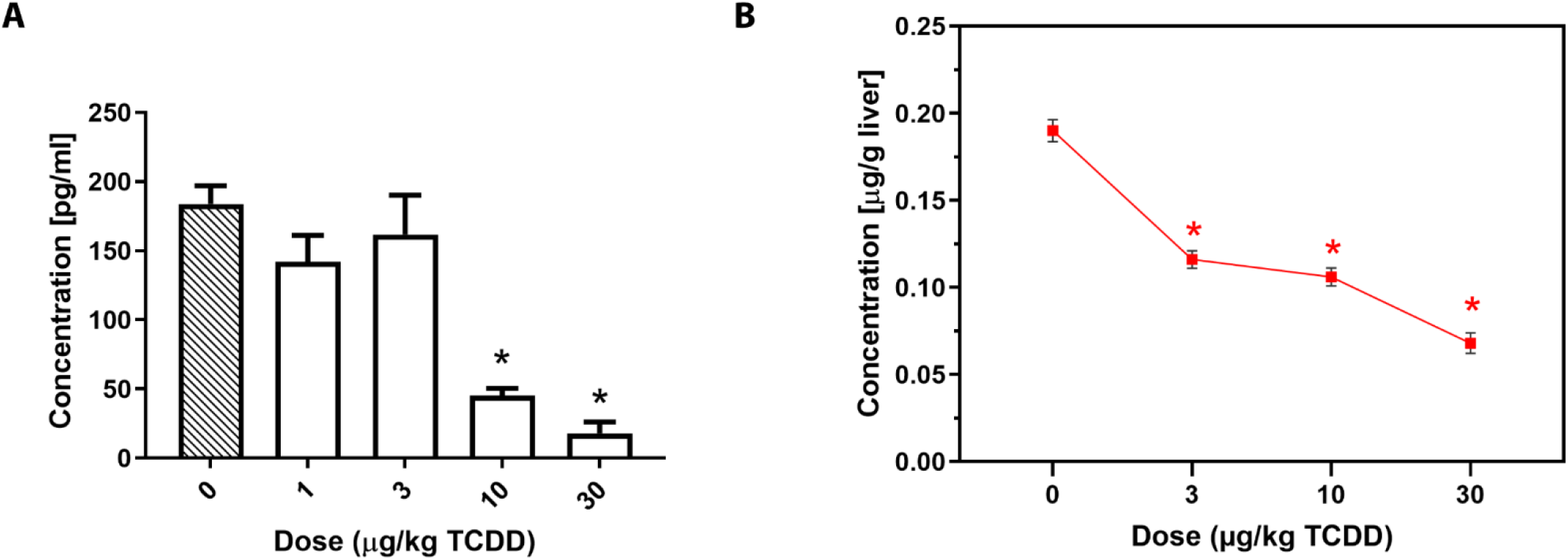
TCDD-elicited effects on **A)** serum cobalamin and **B)** hepatic cobalt levels. Male C57BL/6 mice were orally gavaged every four days with sesame oil vehicle or TCDD for 28 days (n=4-5, ± SEM). Serum cobalamin levels were determined by an ELISA assay. Cobalt levels in liver extracts were measured by inductively coupled plasma mass spectrometry (ICP-MS). Asterisk (*) denotes p<0.05 determined by one-way ANOVA with a Dunnett’s *post-hoc* test.

### Effects on intestinal Cbl absorption and transport

We next examined the effects of TCDD on the expression of genes associated with intestinal absorption and transport of Cbl. Intrinsic factor (IF, *Cblif*), a glycoprotein required for intestinal Cbl absorption, is secreted by parietal cells of the gastric mucosa and therefore was not examined in this study. Cubilin (CUBN) located on the brush border of enterocytes facilitates the endocytic uptake of IF-Cbl complexes. Appreciable levels of *Cubn* expression were detected in duodenal, jejunal, ileal and colonic intestinal segments (**Fig. 4**). *Cubn* was dose-dependently repressed in the duodenum, jejunum, proximal ileum and colon (4.2-, 16.7-, 4.6- and 2.0-fold, respectively), but induced 1.9-fold in the distal ileum. Cbl is then released into the portal circulation in complex with transcobalamin II (TCN2). Repression of *Cubn* in the duodenum, jejunum, proximal ileum and colon segments suggests intestinal Cbl absorption may be inhibited by TCDD. However, the distal ileum is considered the intestinal segment with the greatest Cbl uptake activity (57).

**Figure 4.**
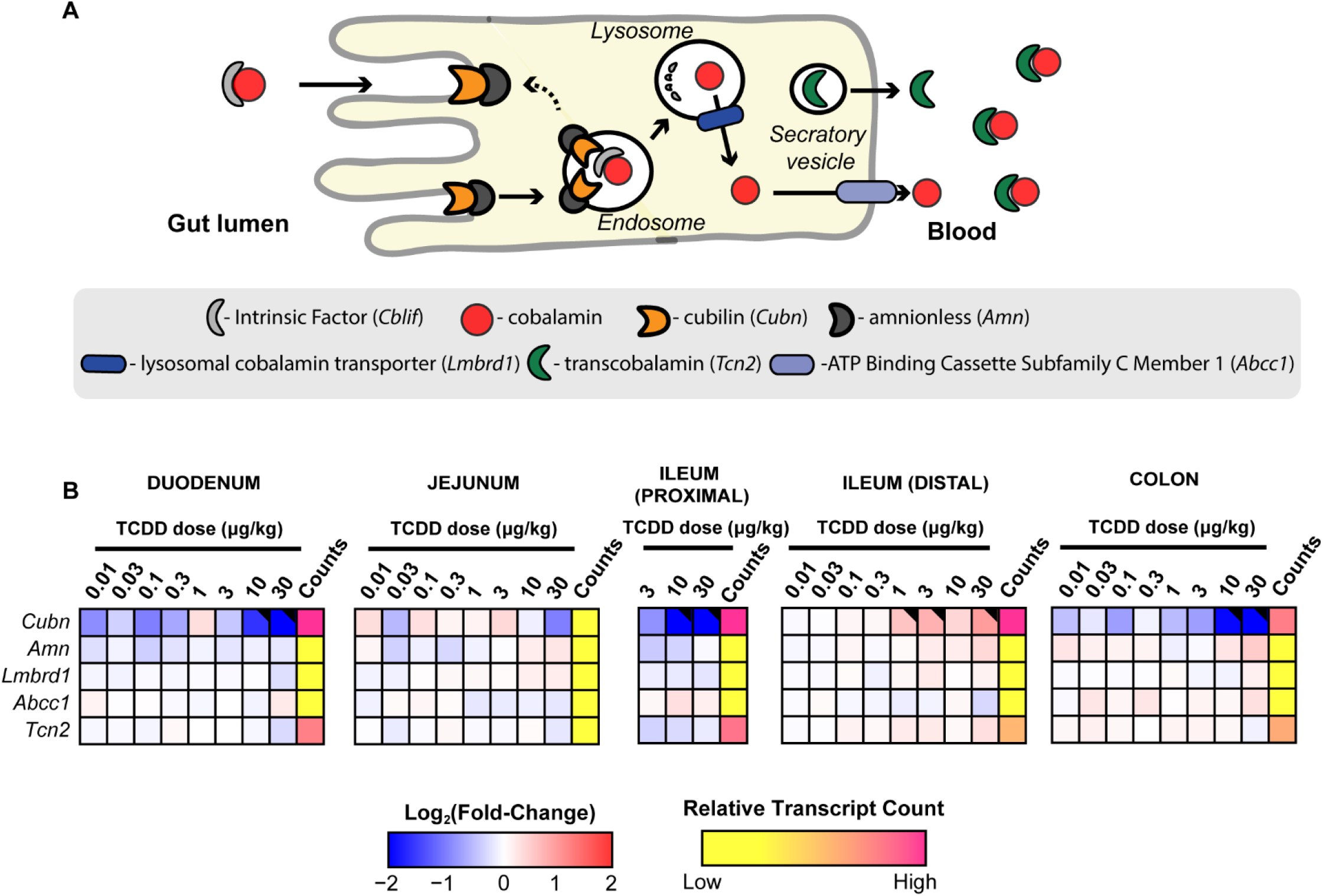
TCDD-elicited effects on gene expression associated with the intestinal absorption and processing of cobalamin (Cbl). **A)** Schematic overview of enterocyte uptake and processing of Cbl. **B)** Dose-dependent effects of TCDD on duodenal, jejunum, ileal (proximal and distal), and colonic gene expression associated with Cbl absorption and processing. Male C57BL/6 mice (n=3) were orally gavaged with sesame oil vehicle or TCDD every 4 days for 28 days. Color scale represents the log2(fold change) for differential gene expression determined by RNA-Seq analysis. Counts represents the maximum raw number of aligned reads to each transcript where a lower level of expression (≤500 reads) is depicted in yellow and a higher level of expression (≥10,000) is depicted in pink. Differential expression with a posterior probability (P1(*t*))>0.80 is indicated with a black triangle in the upper right tile corner.

Cbl deficiency has also been reported following modulation of *de novo* biosynthesis in the gut microbiome and alternatively due to bacterial overgrowth (58, 59). Most gut microbiome taxa possess genes encoding Cbl metabolism associated enzymes. Previous studies have shown that TCDD can alter the gut microbiome, induce bacterial overgrowth and reduce intestinal transit time (14). Metagenomic analyses of cecal contents from this study were assessed to investigate changes in Cbl metabolism by the gut microbiome (14). Gene abundance associated with Cbl biosynthesis and utilization appeared unaffected except for a modest 1.3-fold repression of precorrin-3 methylase (EC 2.1.1.133), an intermediate step in corrin ring biosynthesis, and a 3-fold increase in ABC cobalt transporters (PFAM: PF09819) (**Suppl. Fig. 1 and 2**). Likewise, TCDD had negligible effects on gut microbial propionate metabolism (**Suppl. Table S4**). Based on metagenomics analysis of cecal contents, TCDD elicited negligible effects on microbial Cbl and propionate metabolism.

### Effects on hepatic Cbl uptake, metabolism and trafficking

We also examined the effects of TCDD on gene expression associated with hepatic Cbl uptake, metabolism and trafficking. In humans, circulating Cbl is associated with transcobalamin II (TCN2) or haptocorrin (TCN1) and internalized by hepatocytes following interaction with the transcobalamin receptor (CD320) or the asialoglycoprotein (ASGR1 and 2), respectively, for delivery to lysosomes where TCN2 is degraded and Cbl is released (60). *Tcn2* was repressed 1.5-fold but only at 30 μg/kg TCDD. TCN1 is not expressed in mice, and therefore the 2.4- and 3.9-fold repression of *Asgr1* and *2*, respectively, is irrelevant in regards to hepatic Cbl uptake (60, 61). Released Cbl is exported from the lysosomes via the Lysosome Membrane Chaperone 1 (LMBD1) (62) where it binds to methylmalonic aciduria type C and homocystinuria (MMACHC) proteins (60) and then shuttled to cytoplasmic methionine synthase (MTR) and mitochondrial MUT (**Fig. 5**). Additional proteins including methylmalonic aciduria type A (MMAA) and B (MMAB) convert Cbl to the active AdoCbl cofactor required for MUT activity. Overall, TCDD elicited minimal effects on gene expression suggesting hepatic uptake, metabolism and trafficking are not responsible for lower Cbl levels except for *Mmab* which was repressed 2.4-fold at 30 μg/kg.

**Figure 5.**
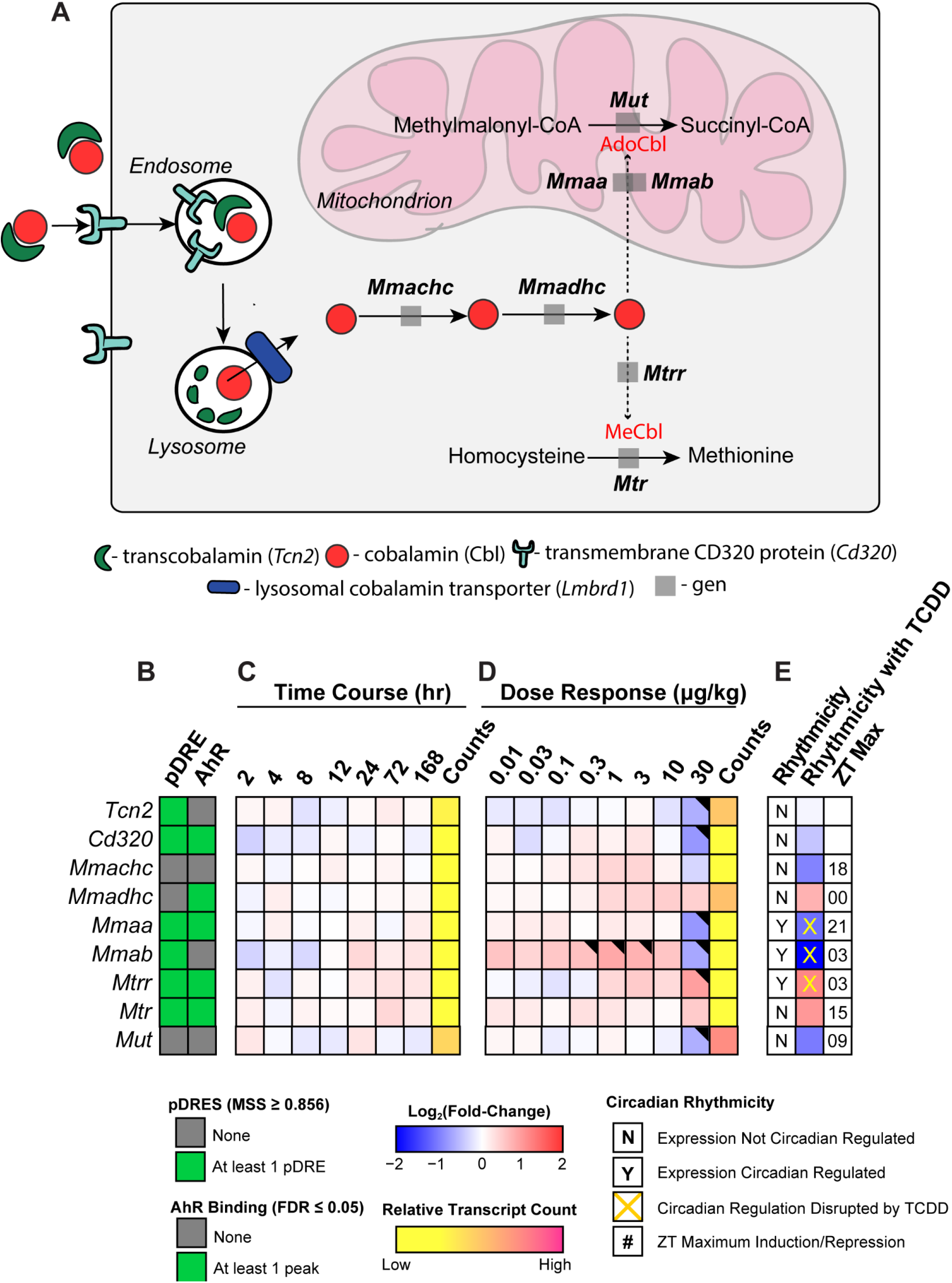
TCDD-elicited effects on gene expression involved in hepatic uptake, metabolism and trafficking of cobalamin (Cbl). **A)** Overview of Cbl uptake, metabolism and trafficking in the mouse liver. **B)** The presence of putative dioxin response elements (pDREs) and AhR genomic binding 2 hrs after a single bolus dose of 30 μg/kg TCDD. **C)** Hepatic expression of genes associated with Cbl uptake, metabolism and trafficking in a time course study. Male C57BL/6 mice (n=3) were administered a single bolus dose of 30 μg/kg TCDD. Liver samples were collected at the corresponding time point. Color scale represents the log2(fold change) for differential gene expression determined by RNA-Seq analysis. Counts represent the maximum number of raw aligned reads for any treatment group. Low counts (<500 reads) are denoted in yellow with high counts (>10,000) in pink. **D)** Dose-dependent gene expression was assessed in mice (n=3) following oral gavage with sesame oil vehicle or TCDD. **E)** Circadian regulated genes are denoted with a “Y”. An orange ‘X’ indicates disrupted diurnal rhythm following oral gavage with 30 μg/kg TCDD every 4 days for 28 days. ZT indicates statistically significant (P1(*t*)>0.8) time of maximum gene induction/repression. Differential expression with a posterior probability (P1(*t*))>0.80 is indicated with a black triangle in the upper right tile corner.

### Cbl depletion and MUT inhibition

Targeted metabolomics analysis of hepatic extracts detected a dose dependent increase in itaconic acid, an immunomodulatory and antimicrobial metabolite of aconitate produced by aconitate decarboxylase 1 (ACOD1 aka IRG1) in macrophages (63) (**Fig. 6M**). Furthermore, hepatic *Acod1* exhibited time-dependent induction following oral gavage with 30 μg/kg TCDD in the absence of AhR genomic enrichment at 2 hrs (**Fig. 6B)**. The time-dependent induction of *Acod1* coincided not only with increased ALT levels (64) but also the time-dependent infiltration of immune cells as indicated by the increased expression of macrophages markers, *Adgre1, Cd5l*, and *Csfr1* (2.3-, 1.9-, and 1.9-fold, respectively) after a single bolus oral gavage of TCDD (**Fig. 6B, K**). *Adgre1* and *Cd5l* did not exhibit AhR genomic enrichment at 2 hrs suggesting increases were due to macrophages recruitment and/or proliferation while *Csf1r* induction may involve the AhR. The low number of macrophages within control livers precludes distinguishing *Adgre1, Cd5l*, and *Csfr1* increases due to induction by TCDD from hepatic macrophage infiltration and/or hepatic macrophages proliferation (65). TCDD also dose-dependently induced *Acod1* (**Fig. 6C**). Discrepancies in the fold-changes between the time course, dose response and circadian studies are consistent with the erratic rhythmic expression of *Acod1* over the 24 hr time period (**Fig 6D**) (14). Itaconate can be activated to itaconyl-CoA which can then interact with the 5-deoxyadenosyl moiety of AdoCbl to form an uncharacterized adduct that disrupts auxiliary repair protein interactions, inactivates AdoCbl, and reduces Cbl levels that inhibit MUT activity (66, 67). These finding are in agreement with the present study where increased itaconic acid levels coincided with diminished Cbl level and inhibited MUT activity. Collectively, the data suggest itaconate was produced following *Acod1* in macrophages that were activated by TCDD, and could factor in the inhibition of MUT due the formation of the uncharacterized inactive itaconyl-CoA:AdoCbl adduct that would reduce available AdoCbl required for MUT activity.

**Figure 6.**
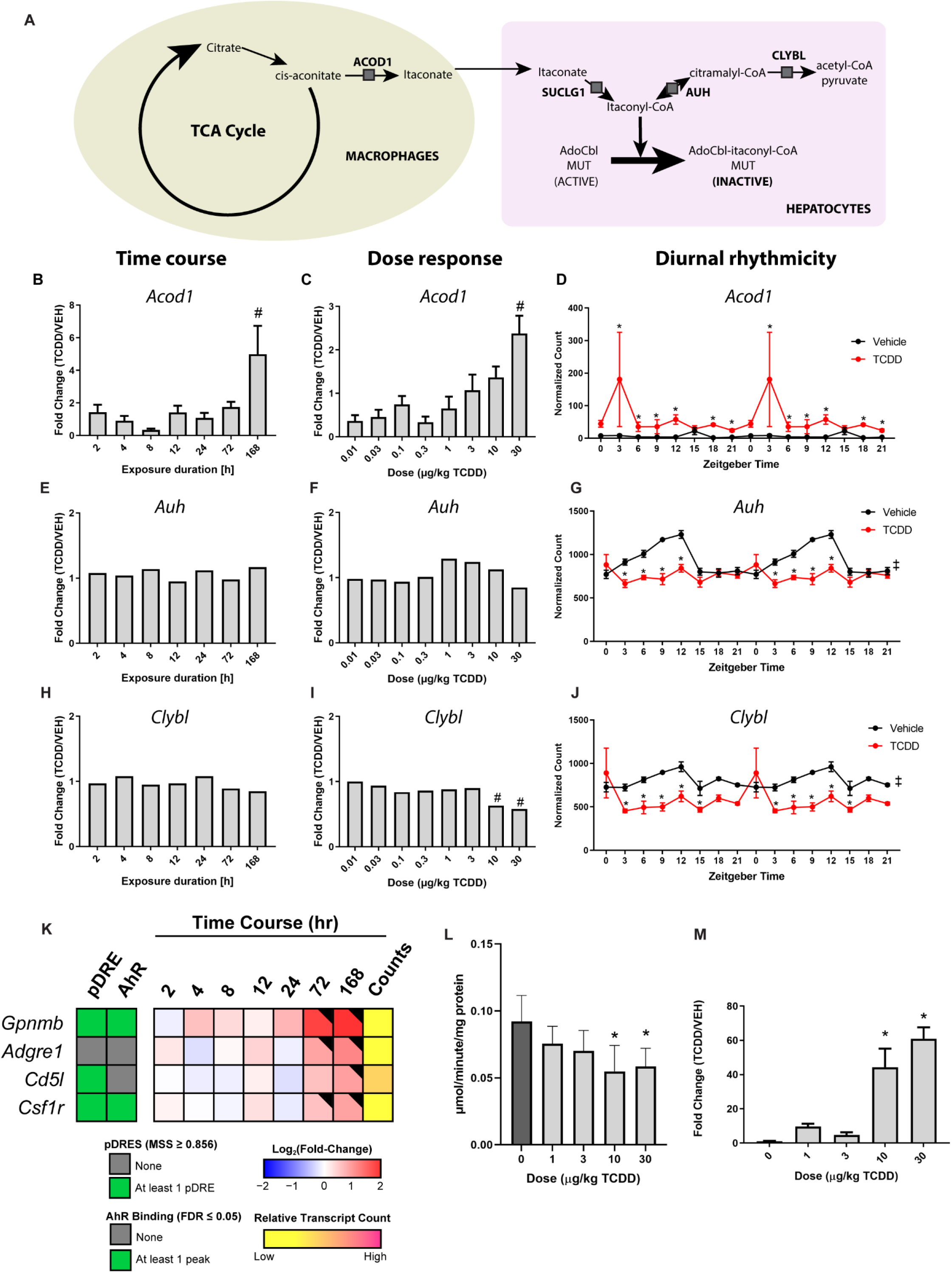
TCDD-elicited effects on the genes involved in the itaconate pathway. **A)** Schematic overview of the pathway. Hepatic expression of **(B)** *Acod1*, **(E)** *Auh*, **(H)** *Clybl* was determined by qRT-PCR (*Acod1*) or RNA-Seq (*Auh, Clybl*) for time-course analysis after a single bolus oral gavage of 30 μg/kg TCDD (n=3-5). Dose dependent expression of **(C)** *Acod1*, **(F)** *Auh*, **(I)** *Clybl* determined by qRT-PCR (*Acod1*) or RNA-Seq (*Auh, Clybl*) following treated with TCDD. Male C57BL/6 mice (n=3-4) were orally gavaged with sesame oil vehicle or TCDD (0.01–30 μg/kg) every 4 days for 28 days. Pound sign (#) denotes p<0.05 determined by one-way ANOVA with a Dunnett’s post-hoc test. The effect of TCDD on the diurnal rhythmicity of (**D**) *Acod1* **(G)** *Auh*, **(J)** *Clybl* in male C57BL/6 mice following oral gavage with sesame oil vehicle or 30 μg/kg TCDD every 4 days for 28 days. Posterior probabilities (*P1(t)≥0.80) comparing vehicle and TCDD were determined using an empirical Bayes method. Diurnal rhythmicity was assessed using JTK_CYCLE (‡ indicates q≤0.1). Data are double-plotted along the x-axis to better visualize rhythmic pattern. **K)** Time dependent hepatic expression of macrophages markers. Male C57BL/6 mice (n=3) were administered a single bolus dose of 30 μg/kg TCDD. Liver samples were collected at the corresponding time point. The presence of putative dioxin response elements (pDREs) and AhR enrichment are represented as green boxes. Color scale represents the log2(fold change) for differential gene expression determined by RNA-Seq analysis. Counts represents the maximum raw number of aligned reads to each transcript where a lower level of expression (≤500 reads) is depicted in yellow and a higher level of expression (≥10,000) is depicted in pink. Differential expression with a posterior probability (P1(*t*))>0.80 is indicated with a black triangle in the upper right tile corner. **L)** Effects of TCDD on MUT activity. MUT activity was measure by thiokinase-coupled spectrophotometric assay where the product, succinyl-CoA, is converted by a second enzyme, thiokinase, to succinate and CoA. Formation of CoA was monitored by the thiol-sensitive reagent, DTNB, which forms a mixed disulfide with CoA and the nitrobenzene thiolate anion (TNB). Results are expressed as μmol TNB formed/minute/mg of protein. Asterisk (*) denotes p < 0.05 determined by one-way ANOVA with a Dunnett’s post-hoc test (n=6). **M)** itaconic acid in liver extracts from male mice orally gavaged every 4 days for 28 days with sesame oil vehicle or TCDD measured by LC-MS. Asterisk (*) denotes p < 0.05 determined by one-way ANOVA with a Dunnett’s *post-hoc* test (n=4). Error bars represent + standard error of the mean.

## DISCUSSION

In this study, metabolomics and gene expression datasets were integrated to examine the increase in the levels of SCEC, an acrylyl-CoA conjugate detected in hepatic extracts from mice orally gavaged with TCDD every 4 days for 28 days. Acrylyl-CoA is a highly reactive intermediate that spontaneously reacts with free sulfhydryl groups, including the sulfhydryl group of cysteine to form the SCEC conjugate, an indicator of acrylyl-CoA, a metabolite normally not detected in healthy individuals at appreciable levels (52, 68). Acrylyl-CoA and 3-HP are intermediates in the catabolism of propionyl-CoA to acetyl-CoA and pyruvate via the Cbl-independent β–oxidation-like pathway, and used as biomarkers of inborn metabolic disorders associated with propionic- and methylmalonic acidemia (69) (70). The SCEC conjugate has also been proposed as a biomarker for Leigh syndrome suggesting a deficiency in ECHS1 activity due to acrylyl-CoA accumulation (71). This led us to investigate TCDD-elicited metabolic reprogramming that redirected propionate metabolism from the canonical Cbl-dependent carboxylation pathway that produces succinyl-CoA to the alternative Cbl-independent β-oxidation-like pathway.

Propionyl-CoA is a byproduct of several reactions including the oxidative metabolism of odd numbered carbon fatty acids as well as the catabolism of several amino acids (i.e., methionine, threonine, isoleucine and valine). However, the most likely source of hepatic propionyl-CoA following TCDD treatment is the shortening of C27-bile acid intermediates to mature C24-bile acids (72) since TCDD dose-dependently increased total bile acids in the liver and serum (14). In addition, TCDD inhibits fatty acid oxidation (15, 16, 73), and had negligible effects on gene expression associated with propionate biosynthesis by the gut microbiome. Acyl-CoA dehydrogenases have extremely low activity towards propionyl-CoA as a substrate. Therefore, propionyl-CoA is preferentially metabolized by the Cbl-dependent carboxylation pathway where it is first carboxylated by propionyl-CoA carboxylase (PCC) to (S)-methylmalonyl-CoA and then epimerized to (R)-methylmalonyl-CoA by methylmalonyl-CoA epimerase (MCEE). Finally, the (R)-methylmalonyl-CoA intermediate undergoes rearrangement by Cbl-dependent MUT to produce the anaplerotic intermediate, succinyl-CoA. Expression of *Pcca*, *Pccb, Echs1, Adhfe1* and *Aldh6a1* were repressed by 30 μg/kg TCDD that correlated with increased SCEC levels in hepatic extracts.

MUT is one of two mammalian enzymes that uses a Cbl derivative as a cofactor, the other being MTR. Specifically, MUT requires AdoCbl for the rearrangement of (R)-methylmalonyl-CoA to succinyl-CoA while MTR uses methylcobalamin (MeCbl) to produce methionine from homocysteine (74). Cbl is considered a rare cofactor with levels ranging between 30-700 nM in humans with deficiency caused by inadequate intake, malabsorption, chemical inactivation or inherited disruption of transport or cellular metabolism (75, 76). It is only synthesized by microorganisms with absorption from animal food sources limited to the distal ileum in humans (57). Given the low levels of Cbl and its potential reactivity in three biologically relevant oxidation states, a complex escort system comprising transporters and chaperones has evolved to ensure delivery to mitochondrial MUT and cytosolic MTR. At least nine proteins are dedicated to the absorption, transport, assimilation, derivatization and trafficking of Cbl, AdoCbl and MeCbl (74). Inborn metabolic disorders as well as intestinal bacterial overgrowth that reduce Cbl levels or disrupt delivery have been implicated in methylmalonic aciduria and/or hyperhomocysteinemia (59, 77). TCDD dose dependently decreased serum Cbl and hepatic cobalt levels, yet had minimal effects on gene expression associated with Cbl absorption, transport, assimilation, derivatization and trafficking. The lone exception was *Cubn*, the membrane receptor responsible for the endocytic uptake of IF-Cbl complexes expressed at the apical pole of enterocytes. *Cubn* in the duodenum, jejunum, proximal ileum and colon was dose-dependently repressed between 3 and 30 μg/kg TCDD coinciding with the dose-dependent decrease in hepatic Cbl levels. However, Cbl absorption in humans is primarily attributed to the distal ileum (57). Moreover, the mouse distal ileum exhibited the highest *Cubn* expression levels and was induced by TCDD in the distal ileum compared to the other intestinal segments. Although enticing to suggest TCDD repression of *Cubn* in the duodenum, jejunum, proximal ileum and colon was responsible for the dose-dependent decrease in hepatic Cbl levels, the induction and overall higher basal expression levels of *Cubn* in the ileum, the primary site of Cbl absorption, implies otherwise. *Mmab* was also repressed but only at 30 μg/kg TCDD.

Other potential mechanisms for lowering hepatic Cbl levels also warrant consideration including the effects of TCDD on the intestinal absorption of Cbl. Specifically, Cbl malabsorption has been attributed to intestinal bacterial overgrowth and decreased gastric acid secretion (57, 75). Parietal cells of the gastric mucosa secrete gastric acid that frees Cbl from ingested proteins. In addition, the lower pH of the stomach favors the protective binding of Cbl to salivary haptocorrin (previously known as R binder). Parietal cells also express IF, the glycoprotein responsible for binding Cbl in the higher pH of the small intestine that facilitate absorption by the ileum (75). Commonly prescribed protein pump inhibitors and histamine 2 receptor antagonists have been shown to suppress gastric acid production. Moreover, a large population case-controlled study reported a dose dependent relationship between Cbl deficiency and the use of acid-suppressing prescription medication for 2 or more years (75, 78). The association was more significant with longer duration of use and diminished after treatment was discontinued (78). Similarly, TCDD is reported to decrease gastric acid secretion by reducing secretory volume, acidity and total acid output (79). Collectively, this suggests that prolonged treatment with TCDD may reduce systemic levels of Cbl due to decreased gastric acid secretion.

More recently, lower Cbl levels have been linked to itaconate, a *cis*-aconitate metabolite produced in large quantities by activated macrophages (66, 67). Itaconate possesses anti-inflammatory properties that block pro-inflammatory cytokine release, inhibit reactive oxygen species (ROS) production, activate the master antioxidant regulator NRF2, and induce the anti-inflammatory transcription factor, ATF3 (80). The induction of *Acod1*, which converts *cis*-aconitate to itaconate, is in agreement with the increased expression of macrophage markers, the dose-dependent decrease in hepatic Cbl levels and the increased levels of itaconic acid in hepatic extracts. ACOD1 is also transcriptionally and post-transcriptionally regulated in response to lipopolysaccharide (LPS) and interferon gamma (IFNγ) (80, 81). Consequently, the bacterial overgrowth and leaky gut caused by TCDD (14) suggest LPS induction of *Acod1* given the absence of AhR enrichment. However, the ability of macrophage secreted itaconate to be absorbed by adjacent cells and cause intracellular effects is debated (82). Although itaconate can be activated to itaconyl-CoA, and subsequently metabolized to citramalyl-CoA, there may be other sources of these intermediates (66). Recent studies also show itaconyl-CoA can inhibit MUT activity by forming a yet to be characterized stable adduct with the 5’-deoxyadenosyl moiety of AdoCbl that reduces Cbl levels (67). Overall, the dose-dependent repression of *Pcca/b* and *Mmab*, as well as the decrease in Cbl levels and reduced MUT activity are consistent with TCDD redirecting propionate metabolism from the Cbl-dependent carboxylation pathway to the Cbl-independent β-oxidation-like pathway that involves propionyl-CoA metabolism to acrylyl-CoA and 3-HP intermediates.

Interestingly, the dose-dependent inhibition of β-oxidation by TCDD also caused a dose-dependent increase in other enoyl-CoA species including octenoyl-CoA (15). The next step in the metabolism of enoyl-CoAs, including acrylyl-CoA, involves hydration. Trifunctional protein (MTP) is a multi-subunit enzyme that carries out enoyl-CoA hydratase, hydroxyacyl-CoA dehydrogenase, and 3-ketothiolase activities in the β-oxidation of straight chain fatty acids. The enoyl-CoA hydratase alpha subunit of MTP (HADHA) prefers longer chain (C12-16) enoyl substrates with minimal activity towards short chain (C4) enoyl-CoAs (83). In contrast, ECHS1 preferentially hydrates shorter chain enoyl-CoAs such as C3 crotonyl-CoA, and exhibits diminished binding affinity for longer chain enoyl-CoAs (84), but efficiently hydrates acrylyl-CoA to 3-HP-CoA (85). We have shown TCDD increased octenoyl-CoA levels and that octenoyl-CoA inhibited the hydration of crotonyl-CoA, the preferred substrate of ECHS1 (15, 83). This suggests inhibition of ECHS1 activity by accumulating octenoyl-CoA could also result in the accumulation of acrylyl-CoA and subsequently, the SCEC conjugate. Inborn metabolic disorders that compromise ECHS1 activity have been linked to urinary acrylyl-CoA conjugate accumulation in infants with pathology severity increasing following palmitate loading (68). Coincidentally, hepatic β-oxidation occurs predominantly in the portal region, the zone first exhibiting dose dependent lipid accumulation and immune cell infiltration after TCDD treatment (64). The structural similarity to other short chain acyl-CoA species, such as propionyl-CoA and crotonyl-CoA, further suggests acrylyl-CoA may be a substrate for post-translational acylation that could impose detrimental structural, and regulatory consequences on enzymes and histones that affect protein-protein interactions and cellular location in addition to function (86).

In summary, we propose TCDD dose dependently repressed gene expression and enzymatic activity that redirected propionyl-CoA from the preferred Cbl-dependent carboxylation pathway to the Cbl-independent β–oxidation-like pathway resulting in the accumulation of highly reactive acrylyl-CoA, due to inhibition of ECHS1 as evident by the presence of the SCEC conjugate. The accumulation of triacylgycerols, FAs, cholesterol, cholesterol esters and phospholipids is first observed as macro- and micro-steatosis following TCDD treatment (12, 64). Fatty liver in combination with increased ROS levels from induced oxidoreductase activities such as CYP1A1, XDH/XO and AOX1 increase oxidative stress levels and lipotoxicity, resulting in subsequent inflammation. This is accompanied by disruption of enterohepatic circulation that not only increased bile acid levels and its propionyl-CoA byproduct, but also promoted bacteria overgrowth in the gut, reduced intestinal motility, and increased intestinal permeability as well as the levels of serum LPS and cytokine including IFNγ (14, 35). Circulating LPS and IFNγ would worsen hepatic oxidative stress, and induce ACOD1 activity following the activation of infiltrating macrophages (81). Activated macrophages produce millimolar levels of itaconate with extracellular itaconate taken up and converted to itaconyl-CoA, the MUT inhibitor (63). Itaconyl-CoA inhibition of MUT activity and the repression of genes associated with the Cbl-dependent carboxylation pathway collectively redirect propionyl-CoA to the less efficient Cbl-independent β-oxidation-like pathway where propionyl-CoA is first oxidized to acrylyl-CoA. Under normal conditions, acrylyl-CoA would be hydrated to 3-HP but ECHS1 activity is inhibited by accumulating octenoyl-CoA due to the futile cycling of FA oxidation and acyl-CoA hydrolysis resulting in incomplete β-oxidation (15). Highly reactive acrylyl-CoA would react with available sulfhydryl groups disrupting protein structure and function contributing to the hepatotoxicity burden along with other toxic metabolites such as dicarboxylic acids, produced as a result of chronic metabolic reprograming in response to persistent AhR activation by TCDD. Collectively, this is consistent with the multi-hit mechanism proposed for NAFLD where steatosis progresses to steatohepatitis with fibrosis and serves as a risk factor for more complex metabolic diseases including HCC (2). It also suggests that TCDD and related compounds, and NAFLD, may share common non-genotoxic mechanisms that lead to HCC. Nevertheless, additional studies are needed to determine the relevance of AhR-mediated metabolic reprogramming by TCDD and related compounds in human models.

## SUPPORTING INFORMATION

This article contains supporting information.

## ACKOWLEDGEMENTS

This project was supported by the National Institute of Environmental Health Sciences Superfund Research Program [NIEHS SRP P42ES004911] and R01ES029541 to TRZ. TRZ is partially supported by AgBioResearch at Michigan State University. RRF and WJS were supported by NIEHS Multidisciplinary Training in Environmental Toxicology [T32ES007255].

## CONTRIBUTIONS

RRF, RN and TRZ designed the in-life study. RRF, WJS and RN performed the animal work. KO, RRF and RN performed the experiments. RRF developed the LC-MS method used for the untargeted metabolomics analysis and was responsible for the metagenomics analysis. Mass spectrometric analysis of itaconic acid was completed by KO and ALS. WJS and RN completed the RNAseq data analysis. KO and RRF prepared the figures and tables. KO, RRF and TZ wrote the manuscript. All authors edited and reviewed the manuscript.

