## Supplementary Information for "Dioxin-elicited decrease in cobalamin redirects hepatic propionyl-CoA metabolism to the β–oxidation-like pathway resulting in acrylyl-CoA conjugate accumulation"

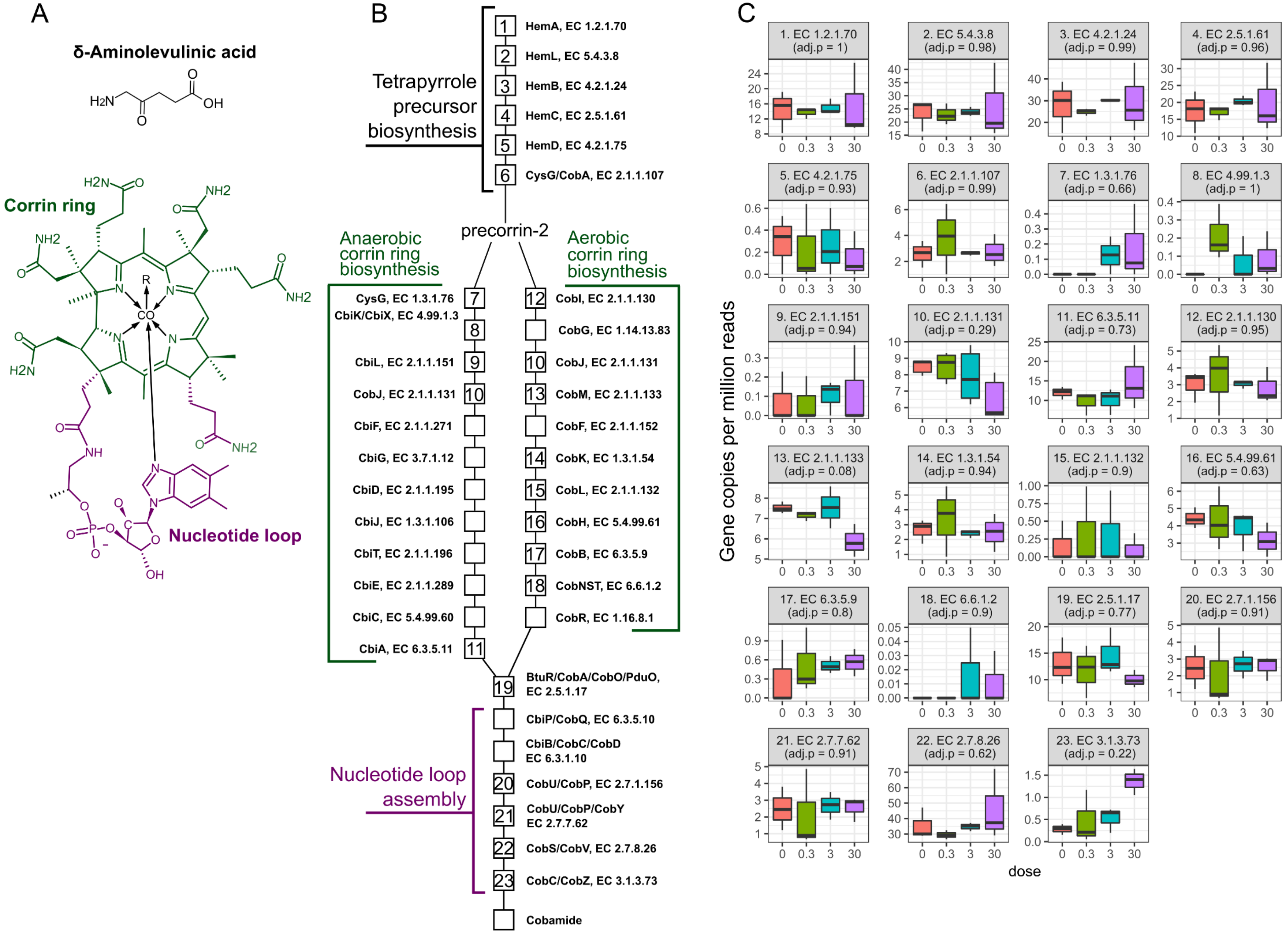
A
δ-Aminolevulinic acid
H2N
O
OH
Corrin ring
H2N
NH2
O
R
Co
N
Nucleotide loop
NH
OH
B
Tetrapyrrole precursor biosynthesis
1 HemA, EC 1.2.1.70
2 HemL, EC 5.4.3.8
3 HemB, EC 4.2.1.24
4 HemC, EC 2.5.1.61
5 HemD, EC 4.2.1.75
6 CysG/CobA, EC 2.1.1.107
precorrin-2
Anaerobic corrin ring biosynthesis
Aerobic corrin ring biosynthesis
7 CysG, EC 1.3.1.76
8 CbiK/CbiX, EC 4.99.1.3
9 CbiL, EC 2.1.1.151
10 CobJ, EC 2.1.1.131
CbiF, EC 2.1.1.271
CbiG, EC 3.7.1.12
CbiD, EC 2.1.1.195
CbiJ, EC 1.3.1.106
CbiT, EC 2.1.1.196
CbiE, EC 2.1.1.289
CbiC, EC 5.4.99.60
11 CbiA, EC 6.3.5.11
12 CobI, EC 2.1.1.130
CobG, EC 1.14.13.83
10 CobJ, EC 2.1.1.131
13 CobM, EC 2.1.1.133
CobF, EC 2.1.1.152
14 CobK, EC 1.3.1.54
15 CobL, EC 2.1.1.132
16 CobH, EC 5.4.99.61
17 CobB, EC 6.3.5.9
18 CobNST, EC 6.6.1.2
CobR, EC 1.16.8.1
19 BtuR/CobA/CobO/PduO, EC 2.5.1.17
Nucleotide loop assembly
CbiP/CobQ, EC 6.3.5.10
CbiB/CobC/CobD EC 6.3.1.10
20 CobU/CobP, EC 2.7.1.156
21 CobU/CobP/CobY EC 2.7.7.62
22 CobS/CobV, EC 2.7.8.26
23 CobC/CobZ, EC 3.1.3.73
Cobamide
C
1. EC 1.2.1.70 (adj.p = 1)
2. EC 5.4.3.8 (adj.p = 0.98)
3. EC 4.2.1.24 (adj.p = 0.99)
4. EC 2.5.1.61 (adj.p = 0.96)
5. EC 4.2.1.75 (adj.p = 0.93)
6. EC 2.1.1.107 (adj.p = 0.99)
7. EC 1.3.1.76 (adj.p = 0.66)
8. EC 4.99.1.3 (adj.p = 1)
9. EC 2.1.1.151 (adj.p = 0.94)
10. EC 2.1.1.131 (adj.p = 0.29)
11. EC 6.3.5.11 (adj.p = 0.73)
12. EC 2.1.1.130 (adj.p = 0.95)
13. EC 2.1.1.133 (adj.p = 0.08)
14. EC 1.3.1.54 (adj.p = 0.94)
15. EC 2.1.1.132 (adj.p = 0.9)
16. EC 5.4.99.61 (adj.p = 0.63)
17. EC 6.3.5.9 (adj.p = 0.8)
18. EC 6.6.1.2 (adj.p = 0.9)
19. EC 2.5.1.17 (adj.p = 0.77)
20. EC 2.7.1.156 (adj.p = 0.91)
21. EC 2.7.7.62 (adj.p = 0.91)
22. EC 2.7.8.26 (adj.p = 0.62)
23. EC 3.1.3.73 (adj.p = 0.22)
Gene copies per million reads
0 0.3 3 30
dose

**Supplementary Figure S1**. Effects of TCDD on microbial Cbl biosynthesis**.** *De novo* Cbl biosynthesis in the gut microbiome involves multiple pathways and >20 enzymes including sub-pathways for δ-aminolevulinic acid (dALA), a tetrapyrrole precursor, the corrin ring and the addition of a nucleotide loop. **(A)** The molecular structure of dALA and Cbl. The corrin ring (green) and nucleotide loop substructures are indicated. **(B)** Schematic depicting Cbl biosynthesis. Each reaction is labeled with an associated gene symbol and enzyme commission (EC) id, and grouped and colored by sub-pathways described in **A**. Genes identified in metagenomic dataset mapping to Cbl biosynthesis EC ids are also denoted by numbers in boxes corresponding to respective graphs in **C**. **(C)** Abundance of genes coding for cbl biosynthesis EC ids assessed by metagenomics analysis of cecum from C57bl/6 mice gavaged with sesame oil vehicle or 0.3, 3, or 30 µg/kg TCDD every 4 days for 28 days. P values adjusted for multiple comparison (adj. p) are denoted in title of respective graphs.

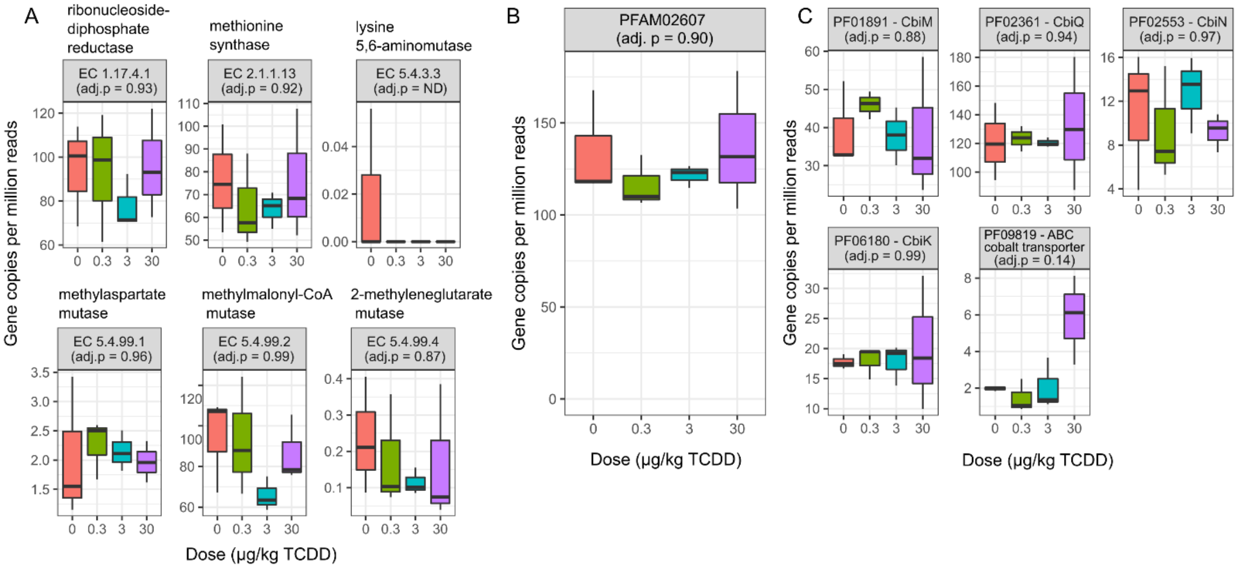


**Supplementary Figure S2**. Effects of TCDD on microbial genes associated with Cbl metabolism**.** Gut microbiome taxa possess genes encoding for Cbl-associated enzymes. Bacterial overgrowth can reduce Cbl bioavailability for the host. The effects bacterial overgrowth induced by TCDD on gut microbiota genes associated with Cbl and cobalt and cobalamin utilization were assessed by metagenomics including **(A)** genes mapping to enzyme commission ids associated with Cbl-dependent enzymes, **(B)** genes containing a PFAM defined Cbl-binding domain (PF02607), and **(C)** genes mapping to PFAMs associated with cbl or cobalt transporters. P values adjusted for multiple comparison (adj. p) are denoted in titles of respective graphs. ND-not determined.

**Supplementary Table S1.** LC-MSMS characteristics of metabolites.

| Analyte | Transition *m/z* | Dwell (sec) | Cove voltage | Collision energy |
|---|---|---|---|---|
| S-(2-carboxyethyl) cysteine | 194.00 > 105.00 | 0.081 | 10 | 16 |
| $^{13}C_5$,$^{15}$N-Methionine | 156.10 > 109.10 | 0.081 | 19 | 9 |

**Supplementary Table S2.** Propionyl-CoA fold changes in comparison to vehicle in liver extracts (n=4-5, ± S.E.) assessed by untargeted liquid chromatography tandem mass spectrometry. Mice were orally gavaged every 4 days for 28 days with TCDD (or sesame oil vehicle). Asterisk (*) denotes significance (p≤0.05) determined by one-way ANOVA with Dunnett's *post-hoc* testing. Score is based on mass error, isotope distribution similar and fragmentation.

| Compound ID | Annotation | Score | Retention Time (min) | Fold-Change (TCDD vs Vehicle) | | | | |
|---|---|---|---|---|---|---|---|---|
| | | | | 0.3 µg/kg | 1 µg/kg | 3 µg/kg | 10 µg/kg | 30 µg/kg |
| HMDB01275 | Propionyl-CoA | 42.4 | 6.84 | 1.56 ± 0.23 | 0.55 ± 0.14 | 0.89 ± 0.16 | 0.11 ± 0.07* | 0.19 ± 0.07* |

**Supplementary Table S3**: QRTPCR Primers. sequences (5' -3') and product sizes for genes analyzed by qRT-PCR.

| Gene ID | Gene Symbol | Gene Name | Primer sequence | | Product length |
|---|---|---|---|---|---|
| | | | Forward | Reverse | |
| 11461 | *Actb* | Actin, beta | GCTACAGCTTCACCACCACA | TCTCCAGGGAGGAAGAGGAT | 123 |
| 14433 | *Gapdh* | glyceraldehyde-3-phosphate dehydrogenase | GTGGACCTCATGGCCTACAT | TGTGAGGGAGATGCTCAGTG | 125 |
| 15452 | *Hprt* | hypoxanthine guanine phosphoribosyl transferase | AAGCCTAAGATGAGCGCAAG | TTACTAGGCAGATGGCCACA | 104 |
| 16365 | *Acod1* | aconitate decarboxylase 1 | TATGCCAACTACTCCCCCGA | CGGGAAGCTCTTAAAGGCCA | 90 |

**Supplementary Table S4. Microbial metabolism of propionate**. Gene abundance (counts per million copies) of reads associating with uniref90 ids and mapping to enzyme commission (EC) entries where propionyl-CoA or propionate are listed as substrates in respective enzymatic reaction. Significance (*) denotes adjusted p (adj. p) < 0.05.

| EC entry | Name | Counts per million copies (CPM) | | | | |
|---|---|---|---|---|---|---|
| | | 0 μg/kg | 0.3 μg/kg | 3 μg/kg | 30 μg/kg | adj. p |
| 2.3.1.16* | Acetyl-CoA C-acyltransferase | 0.15 ± 0.12 | 0.20 ± 0.07 | 0.47 ± 0.48 | 1.59 ± 0.32 | 0.04 |
| 4.1.3.32 | 2,3-dimethylmalate lyase | 0.19 ± 0.06 | 0.15 ± 0.07 | 0.33 ± 0.16 | 0.35 ± 0.10 | 0.50 |
| 1.3.8.1 | Short-chain acyl-CoA dehydrogenase | 16.92 ± 11.97 | 13.33 ± 2.99 | 14.43 ± 3.31 | 9.43 ± 4.72 | 0.61 |
| 1.3.1.95 | Acryloyl-CoA reductase | 0.13 ± 0.12 | 0.13 ± 0.08 | 0.29 ± 0.13 | 0.07 ± 0.06 | 0.67 |
| 2.8.3.1 | Propionate CoA-transferase | 0.15 ± 0.08 | 0.01 ± 0.02 | 0.15 ± 0.05 | 0.03 ± 0.03 | 0.68 |
| 2.8.3.1 | Propionate CoA-transferase | 0.15 ± 0.08 | 0.01 ± 0.02 | 0.15 ± 0.05 | 0.03 ± 0.03 | 0.68 |
| 6.2.1.1 | Acetate--CoA ligase | 0.57 ± 0.33 | 0.74 ± 0.61 | 0.21 ± 0.10 | 0.26 ± 0.22 | 0.68 |
| 6.2.1.17 | Propionate--CoA ligase | 0.00 ± 0.00 | 0.01 ± 0.02 | 0.07 ± 0.12 | 0.05 ± 0.06 | 0.77 |
| 1.2.7.1 | Pyruvate synthase | 9.79 ± 3.19 | 15.04 ± 1.41 | 15.11 ± 4.00 | 10.72 ± 2.31 | 0.84 |
| 2.1.3.1 | Methylmalonyl-CoA carboxytransferase | 8.90 ± 1.23 | 9.01 ± 2.26 | 6.73 ± 0.92 | 7.54 ± 2.22 | 0.84 |
| 1.3.1.84 | Acrylyl-CoA reductase | 0.00 ± 0.00 | 0.00 ± 0.00 | 0.03 ± 0.04 | 0.02 ± 0.04 | 0.86 |
| 6.4.1.3 | Propionyl-CoA carboxylase | 35.63 ± 6.67 | 35.57 ± 7.73 | 28.68 ± 2.48 | 31.16 ± 7.60 | 0.87 |
| 2.7.2.15 | Propionate kinase | 0.00 ± 0.00 | 0.00 ± 0.00 | 0.02 ± 0.04 | 0.01 ± 0.02 | 0.89 |
| 1.2.1.27 | Methylmalonate-semialdehyde dehydrogenase | 0.00 ± 0.00 | 0.03 ± 0.05 | 0.00 ± 0.00 | 0.02 ± 0.04 | 0.91 |
| 2.3.1.8 | Phosphate acetyltransferase | 18.84 ± 4.80 | 17.95 ± 7.31 | 13.69 ± 2.42 | 15.27 ± 2.95 | 0.91 |
| 2.3.3.5 | 2-methylcitrate synthase | 0.15 ± 0.09 | 0.05 ± 0.06 | 0.12 ± 0.13 | 0.19 ± 0.21 | 0.93 |
| 2.3.1.54 | Formate C-acetyltransferase | 4.53 ± 1.95 | 8.42 ± 2.03 | 7.70 ± 5.02 | 7.03 ± 3.14 | 0.96 |
| 2.3.1.222 | Phosphate propanoyltransferase | 29.62 ± 9.16 | 22.04 ± 4.18 | 20.68 ± 5.00 | 29.71 ± 19.89 | 0.99 |
| 2.3.1.9 | Acetyl-CoA C-acetyltransferase | 11.55 ± 7.45 | 10.60 ± 3.19 | 11.74 ± 3.00 | 11.05 ± 3.62 | 0.99 |
| 2.7.2.1 | Acetate kinase | 137.43 ± 7.41 | 141.50 ± 16.41 | 141.03 ± 8.14 | 140.25 ± 6.24 | 0.99 |
